# Epidemic foci of hemorrhagic fever with renal syndrome in Shandong Province, China, based on patients, rodents and molecular epidemiology characteristics, 2012-2015

**DOI:** 10.1101/518274

**Authors:** Zhaolei Zheng, Peizhu Wang, Zhiqiang Wang, Dandan Zhang, Xu Wang, Shuqing Zuo, Xiujun Li

## Abstract

**Background:** Hemorrhagic fever with renal syndrome (HFRS), an infectious disease caused by hantaviruses, is endemic in China and remains a serious public health problem. Historically, Shandong Province has had the largest HFRS burden in China. However, we do not have a comprehensive and clear understanding of the current epidemic foci of HFRS in Shandong Province.

**Methodology/Principal Findings:** The incidence and mortality rates were calculated, and a phylogenetic analysis was performed after laboratory testing of the virus in rodents. Spatial epidemiology analysis was applied to investigate the epidemic foci, including their sources. A total of 6,206 HFRS cases and 59 related deaths were reported in Shandong Province. The virus carriage rates of the rodents *Rattus norvegicus*, *Apodemus agrarius* and *Mus musculus* were 10.24%, 6.31% and 0.27%, respectively. The phylogenetic analysis indicated that two novel viruses isolated from *R. norvegicus* in Anqiu City and Qingzhou City were dissimilar to the other isolated strains, but closely related to strains previously isolated in northeastern China. Three epidemic foci were defined, two of which were derived from the Jining and Linyi epidemic foci, respectively, while the other was the residue of the Jining epidemic focus.

**Conclusions/Significance:** The southeastern and central Shandong Province are current key HFRS epidemic foci dominated by *A. agrarius* and *R. norvegicus*, respectively. Our study could help local departments to strengthen prevention and control measures in key areas to reduce the hazards of HFRS.

**Author summary:** Hemorrhagic fever with renal syndrome (HFRS) is a global infectious disease, which is still a serious public health threat in China today. The reported HFRS cases in Shandong Province accounted for approximate one third of total cases in the whole country. HFRS is a zoonosis mainly caused by Hantaan virus (HTNV) and Seoul virus (SEOV), which natural rodent hosts are *A. agrarius* and *R. norvegicus*, respectively. To explore the current HFRS epidemic foci based on patients, rodents and molecular epidemiology characteristics in Shandong Province, we collected the records of HFRS cases from whole province and the rodents captured in 14 surveillance sites. We found that the epidemic situation of HFRS is quiet different in temporal and spatial distribution. Three epidemic foci were defined based on patients, rodents and molecular epidemiology characteristics. The situation of HFRS epidemic foci in Shandong Province was clear. Our study provides a reference for relevant departments to develop key prevention strategies.

## Introduction

Hemorrhagic fever with renal syndrome (HFRS), a global infectious disease caused by hantaviruses (HV) in the *Bunyaviridae* family, is characterized by fever, hemorrhage, kidney damage and hypotension. All the currently known HV genotypes can cause HFRS, including Hantaan virus (HTNV), Seoul virus (SEOV), Puumala virus (PUUV) and Dobrava/Belgrade virus (DOBV/BGDV)[1,2,3]. In China, most cases are caused by two HV genotypes, HTNV and SEOV, which are mainly carried by the rodents *Apodemus agrarius* and *Rattus norvegicus*, respectively.

According to previous studies, over 90% of all cases of HFRS in the world have occurred in China in recent decades[4,5]. Although the harm caused by HFRS in humans has declined in recent years due to the development of preventive measures and medical capabilities, HFRS is still a serious public health threat in China today. Shandong Province, which has the second largest population among all the provinces in China, has been suffering from a large HFRS burden for several decades. In the past decade, Shandong province remained in the top five of all provinces in China regarding the number of HFRS cases[1].

Changes in the epidemic foci (areas in which some wild animals have a long-term preservation of infectious pathogen) of infectious diseases are caused by many factors, including natural, human and viral host factors[6]. In recent years, Shandong Province developed an *R. norvegicus*-dominated mixed viral genotypes epidemic focus. However, the characteristics of the HFRS epidemic foci in Shandong Province have been constantly changing, and the health threat to humans has been constantly changing as well. Adjustments to policies and practices should be performed in a timely manner. However, although many studies have analyzed the HFRS epidemic situation in Shandong Province[4,6,7], we do not have a comprehensive and clear understanding of the current epidemic foci of HFRS. For this reason, we conducted a systematic analysis of surveillance information on HFRS in Shandong Province from 2012 to 2015 to assess the severity of the situation and determine the new epidemic foci in Shandong Province.

## Methods

### Ethics statement

This study was reviewed and approved by the Ethical Review Board, Science and Technology Supervisory Committee at the Beijing Institute of Microbiology and Epidemiology. The animal work described here adhered to the guidelines of the Animal Subjects Research Review Boards at the Beijing Institute of Microbiology and Epidemiology.

### Study area and data source

The study area is located in the North China Plain, in eastern China (34°22’to 38°23’north latitude, 114°19’ to 122°43’ east longitude, Fig 1), with a population of more than 99 million and an area of 158,000 km^2^. Shandong Province has a large population of agricultural workers and high land cultivation rate in China. The central region is mountainous and hilly, which is surrounded by plains.

**Fig 1.**
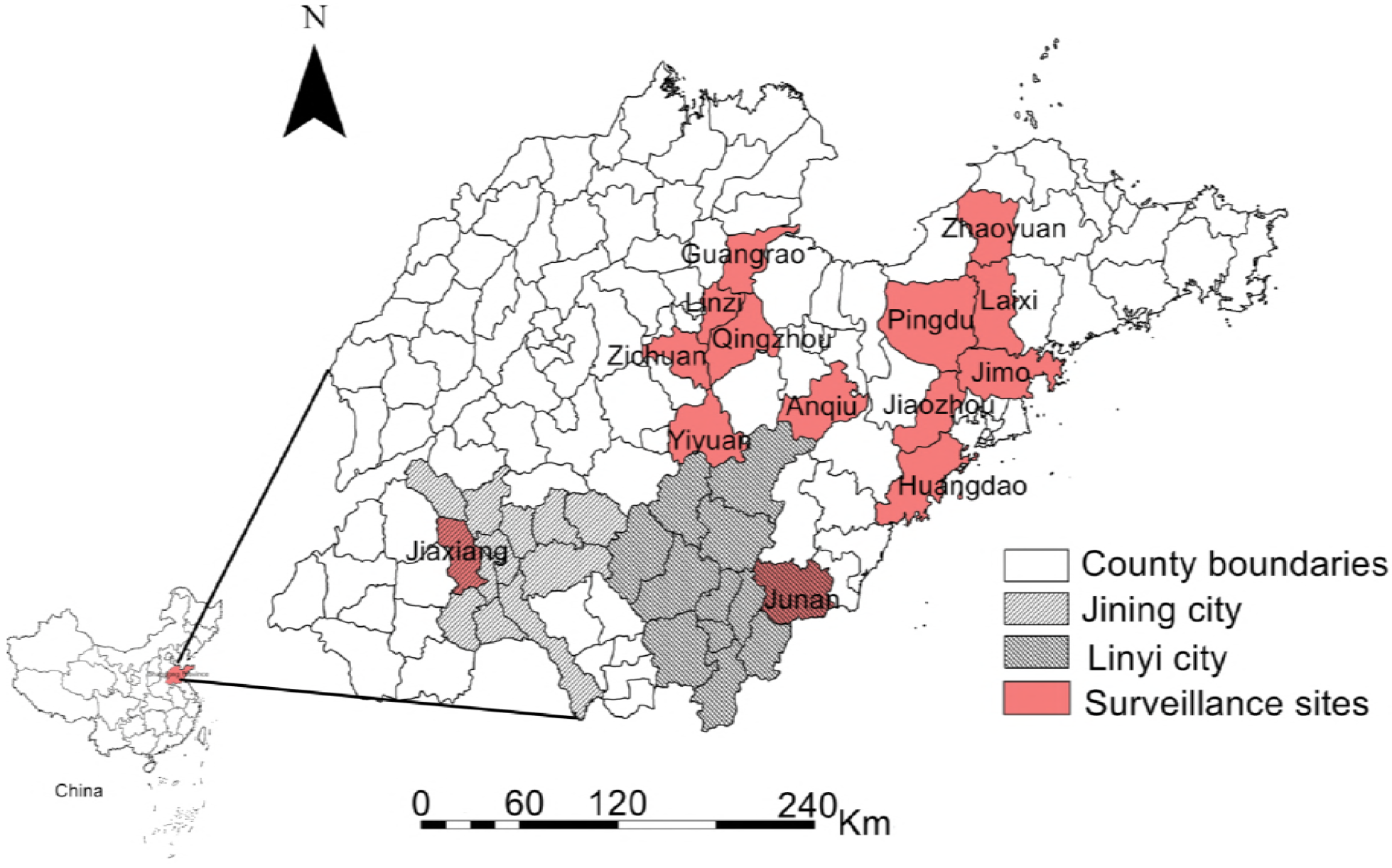
Location of this study. The areas marked by light red are the 14 surveillance counties of this study. The black slash and black mesh line represent the cities where the previous hemorrhagic fever with renal syndrome (HFRS) epidemic foci located in Shandong Province. The map was visualized with ArcGIS 10.2 software (ESRI Inc., Redlands, CA, USA).

The records of HFRS cases between 2012 to 2015 were obtained from the China Information System for Disease Control and Prevention and all data analyzed were anonymized. Based on historical HFRS surveillance data from Shandong Province, 14 representative counties with serious epidemics were selected as the surveillance sites in this study. Rodents surveillance data were also collected from the China Information System for Disease Control and Prevention. Lung samples of the captured rodents were obtained for laboratory testing.

### Epidemiological description

The 2012-2015 records of HFRS cases covering the whole of Shandong Province were used to calculate the monthly incidence rate (per 100,000 population). The proportions of cases in different demographic groups, based on gender, age and occupation were calculated. To observe the temporal distribution of cases more intuitively, we divided all cases into three groups (Feb to May, June to Sept and Oct to Jan) based on the two annual HFRS incidence peaks.

### Rodent surveillance analysis

According to the information on the captured rodents, the proportion of each rodent species out of all the rodents captured was calculated. Lung samples of the captured rodents were screened using the reverse transcription-polymerase chain reaction (RT-PCR). Total RNA, which was extracted from 20–50 mg of lung tissue using the TRIzol reagent (Invitrogen, USA), was reverse-transcribed using M-MLV reverse transcriptase (Invitrogen, USA) and the primer P14: 5’-TAGTAGTAGACTCC-3’[8]. The PCR was performed to amplify a partial L sequence, using primer pairs as previously described[9]. Amplification result was sequenced and compared with data from the National Center for Biotechnology Information (NCBI) database using the Basic Local Alignment Search Tool (BLAST) to determine which genotypes they were[10].

### Phylogenetic analysis

Based on the results of pre-experiment, we screened the isolated strains with low homology in the same county every year. Eventually, the partial L segments of 29 isolated strains from the 14 surveillance sites were selected. Additionally, we searched GenBank and downloaded data on 15 relevant HV strains isolated in other regions of China, the Korean Peninsula and Europe. A phylogenetic tree was then constructed, using the neighbor-joining (NJ) method and bootstrap testing (1000 replicates), in MAGE 7.0 software (Pennsylvania State University, PA, USA)[11].

### Epidemic foci analysis

According to the HFRS cases distribution, molecular epidemiology and rodent surveillance data, the epidemic foci of HFRS in Shandong Province during the study period were defined. The Kriging interpolation method was applied to display the extent of the epidemic foci more clearly.

### Data analysis

Data were analyzed using SAS 9.4 (SAS Institute Inc., Cary, NC, USA). All maps were mapped by using a geographical information system (GIS) technique in ArcGIS 10.2 software (ESRI Inc., Redlands, CA, USA). MEGA 7.0 was used to conduct the phylogenetic analysis.

## Results

### Epidemiological characteristics of HFRS

From 2012 to 2015, 6,206 HFRS cases and 59 related deaths were reported in Shandong Province, and the annual incidence rate, mortality rate and case fatality rate ranged from 1.15 to 1.87 per 100,000, 0.01 to 0.02 per 100,000, and 0.80% to 1.07%, respectively (Table 1). During the study period, the annual HFRS incidence rate in Shandong Province peaked in 2013, then decreased year by year, and reached its lowest point in 2015. In addition, there were two incidence peaks each year, the small peak was from April to June and the larger one was from October to January. However, mortality rates and case fatality rates declined year by year. The deaths mainly occurred from September to January (Fig 2). As for spatial distribution, significant dynamic variation was found in that the high-risk areas were concentrated in a single large region in central and southeastern Shandong Province in 2012, which gradually separated into two regions, and then eventually formed two relatively independent high-risk regions in 2015 (Fig 3).

**Table 1.** Epidemics of HFRS in Shandong province, 2012-2015

| Years | No. patients | No. deaths | Incidence rate<br>(per 100,000) | Mortality<br>(per 100,000) | Fatality(%) |
| --- | --- | --- | --- | --- | --- |
| 2012 | 1,683 | 18 | 1.743 | 0.019 | 1.070 |
| 2013 | 1,799 | 18 | 1.867 | 0.019 | 1.000 |
| 2014 | 1,602 | 14 | 1.639 | 0.014 | 0.874 |
| 2015 | 1,122 | 9 | 1.148 | 0.009 | 0.802 |

HFRS= hemorrhagic fever with renal syndrome.

**Fig 2.**
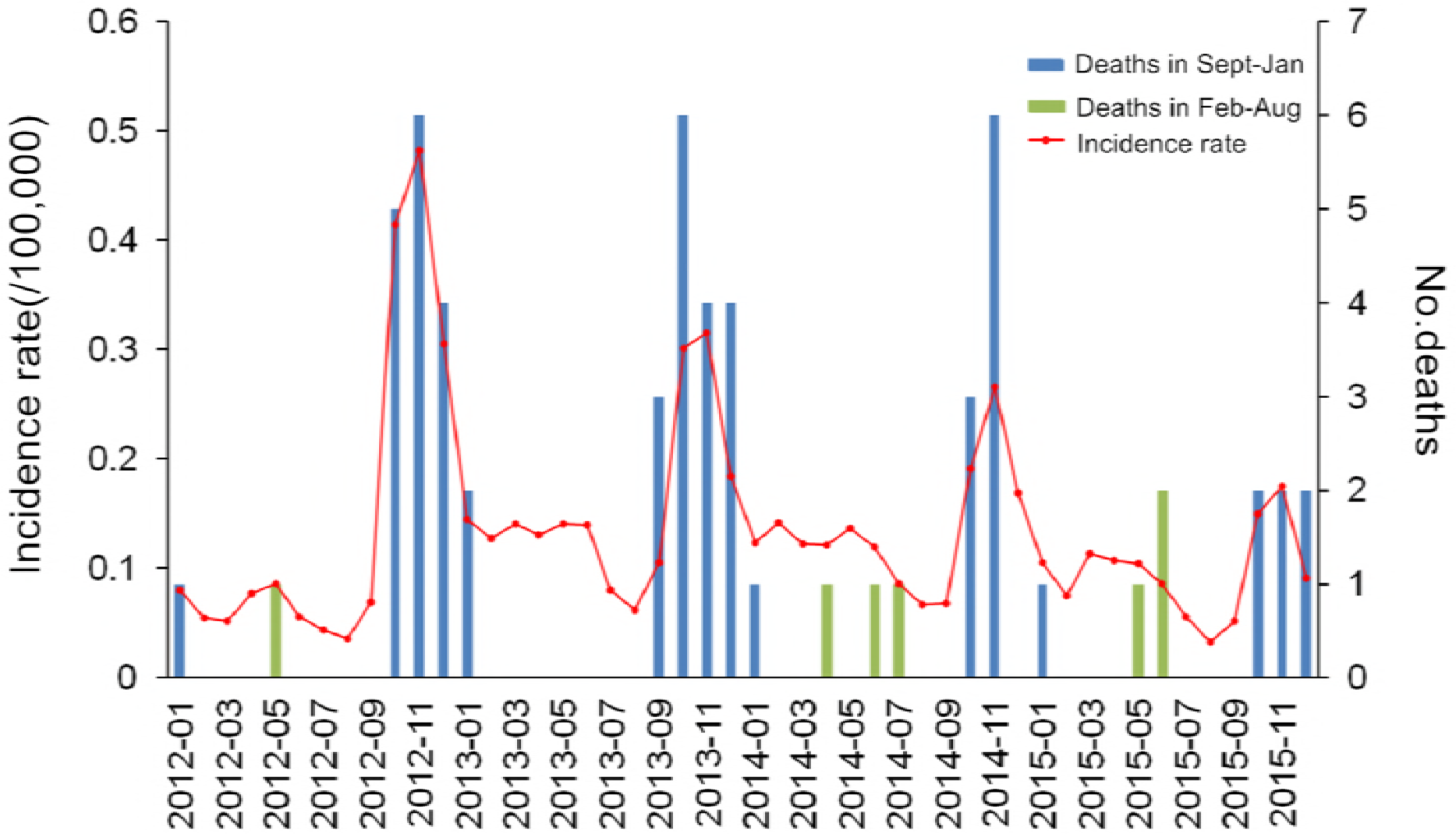
The temporal distribution of hemorrhagic fever with renal syndrome (HFRS) incidence rate and deaths in Shandong Province from 2012 to 2015. The monthly incidence rate and the number of related deaths in Shandong Province were calculated and displayed by curves and bars, respectively. The blue bars represented the deaths occurred between September to January, and green bars represented the deaths occurred in other months.

**Fig 3.**
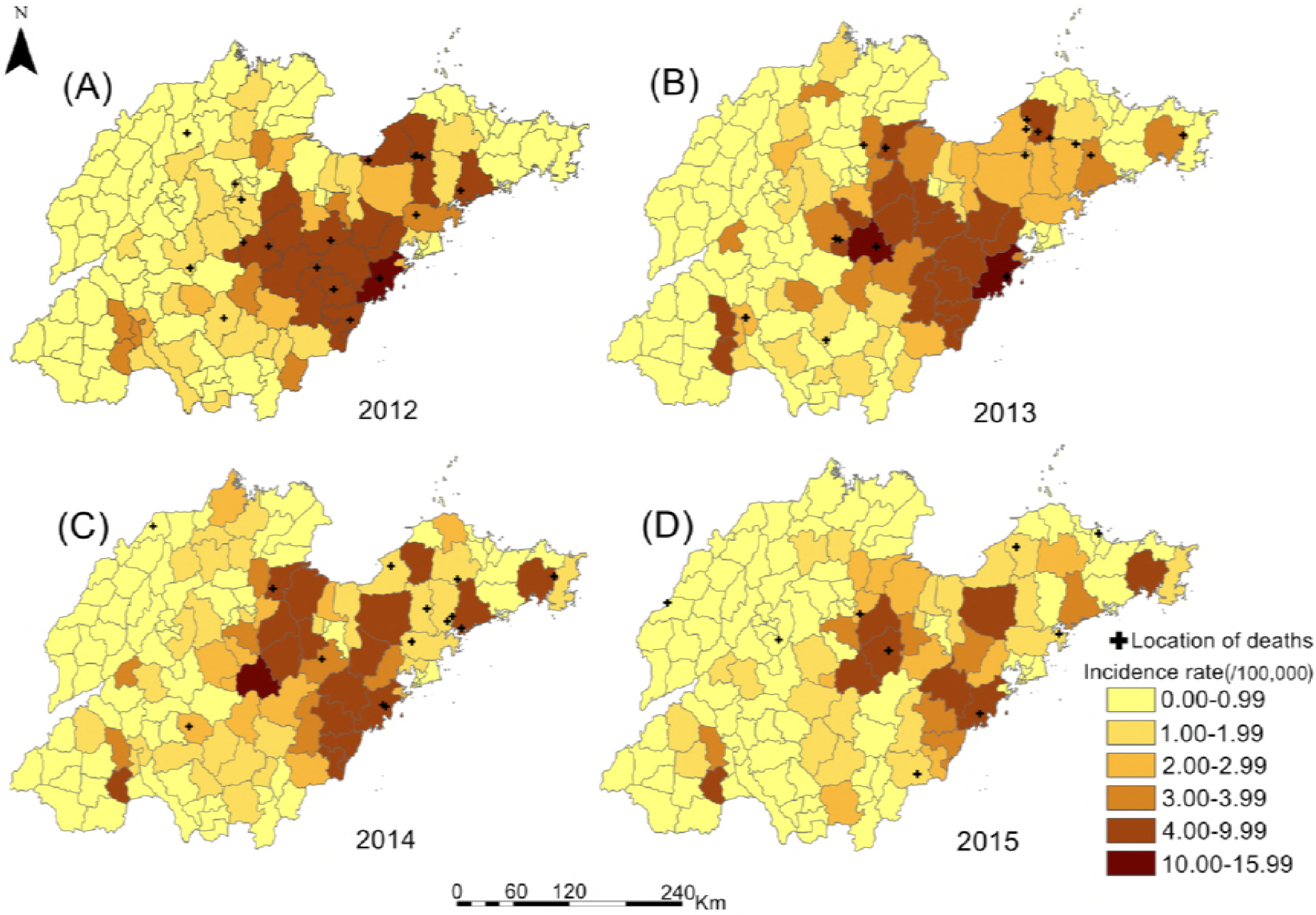
The spatial distribution of hemorrhagic fever with renal syndrome (HFRS) incidence rates and deaths in Shandong province 2012-2015. HFRS incidence rates in 142 counties of Shandong Province were calculated and divided into 6 groups represented by different colors. The plus sign represented the location of deaths. The map was visualized with ArcGIS 10.2 software (ESRI Inc., Redlands, CA, USA).

Regarding the demographic characteristics, 4465 males accounted for 72% of all cases. We found that 81% of all cases occurred in individuals aged 30–70 years, with the 41–50-year-old age group having the highest proportion of all the 10-year age groups. The three occupations with the highest incidence rates were farm workers (85%), urban workers (5%) and students (3%). The results clearly showed that middle-aged male farm workers are the main population to experience HFRS in Shandong Province. (Table 2).

**Table 2.**
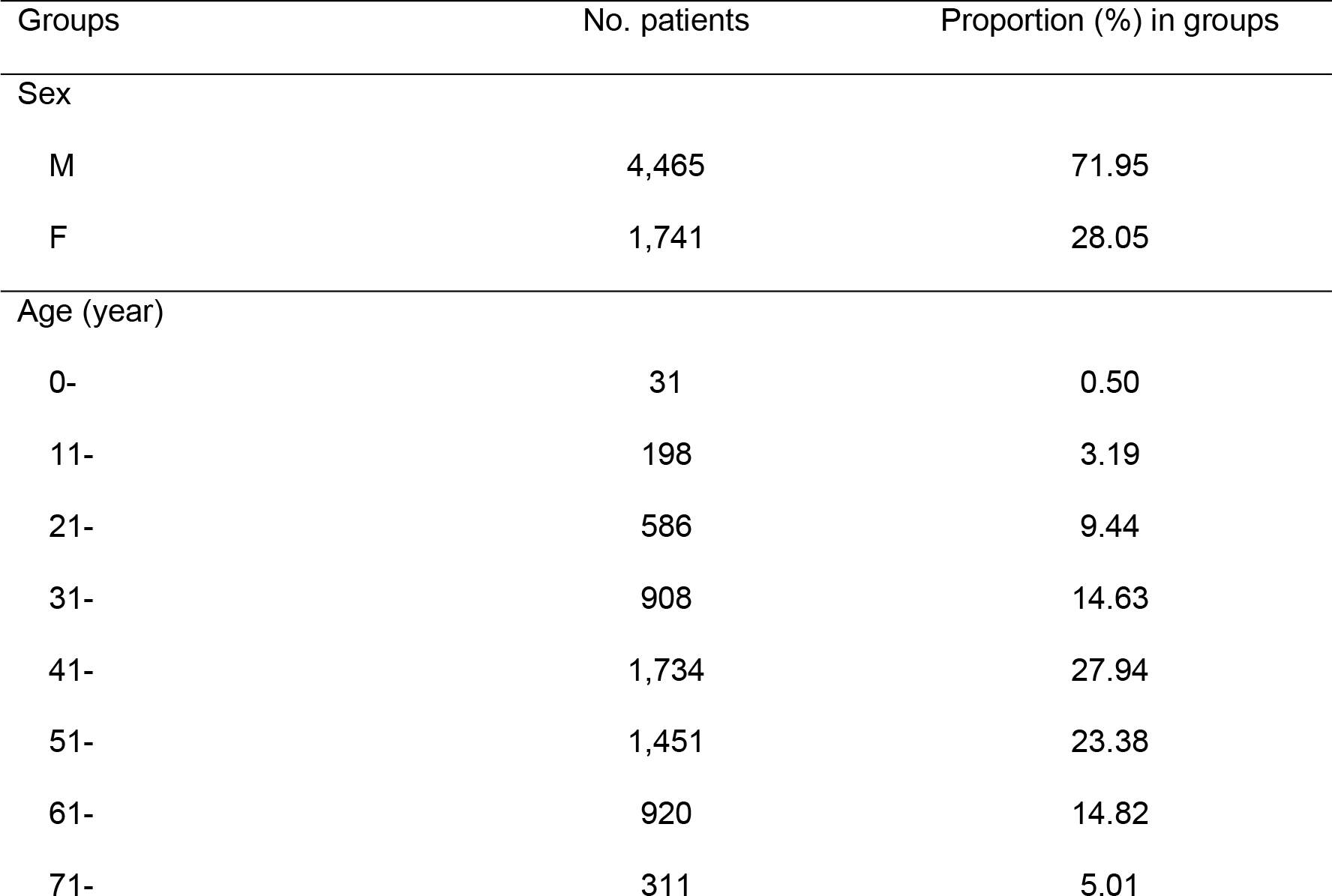
Demographic distribution of HFRS cases in Shandong Province from 2012 to 2015

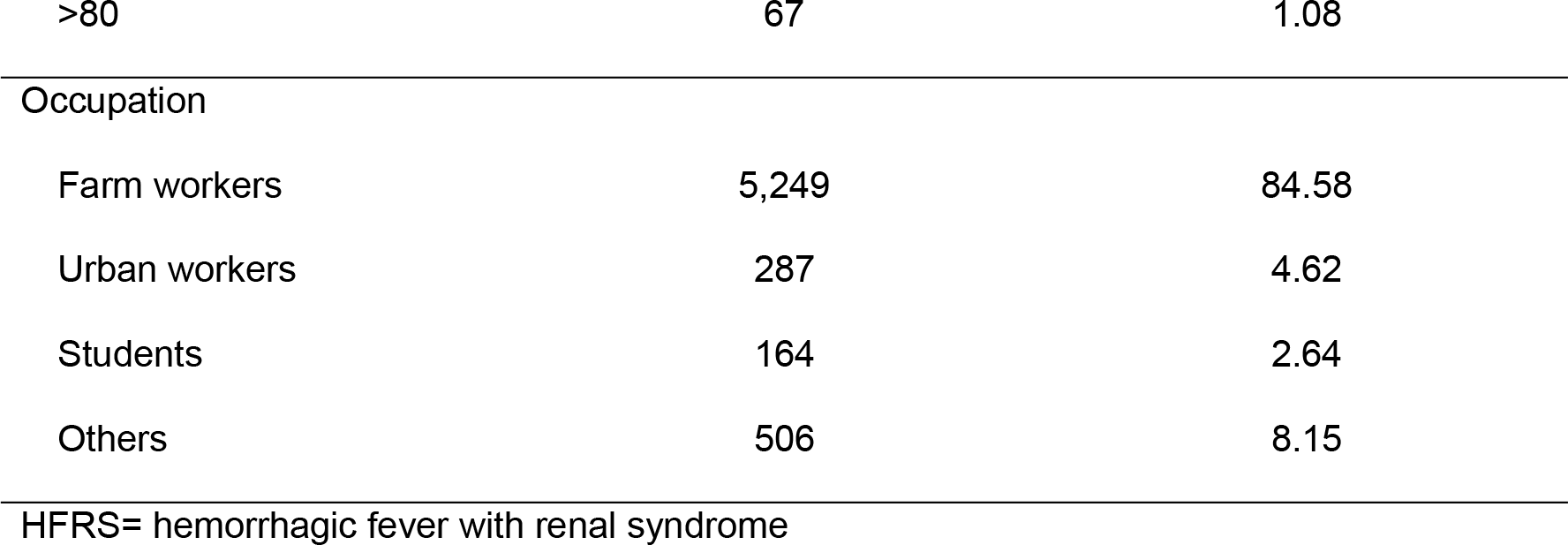

### Surveillance of rodent hosts

A total of 1,798 rodents were captured in 14 rodent surveillance sites in Shandong Province from 2012 to 2015, including 908 *R. norvegicus*, 373 *Mus musculus*, 269 *A. agrarius*, 95 *Cricetulus barabensis*, 73 *Sorex araneus*, 31 *Cricetulus triton*, 25 *Niviventer confucianus*, 23 *Rattus flavipectus* and 1 *Miorotus fortis*. The proportions of different rodent species indicated that the dominant species in the surveillance sites was *R. norvegicus*, followed by *Mus musculus* and *A. agrarius*.

The laboratory testing results revealed that 111 of the captured rodents were positive for HV, of which 93 were *R. norvegicus*, 17 were *A. agrarius* and 1 was *Mus musculus*. The mean HV carriage rate among all rodents was 6.17%, and the rates of *R. norvegicus*, *A. agrarius* and *Mus musculus* were 10.24%, 6.32% and 0.27%, respectively (Table 3). It can be seen that the HV carriage rates in this study were highest for *R. norvegicus* and *A. agrarius* among all the rodent species. However, it is worth noting that one HV strain was also isolated from a single *Mus musculus* in this study.

We constructed a geographic map of the different HV-infected rodent species. This indicated that the HV-infected dominant rodents in central Shandong Province were *R. norvegicus*, followed by *Mus musculus*, while *A. agrarius* was the dominant HV-infected rodent in southeastern Shandong Province (Fig 4A). Additionally, the HV carriage rates of rodents in each surveillance site were calculated and plotted on another map (Fig 4B, S1 Table). This showed that the surveillance sites with high HV carriage rates were mainly concentrated in central Shandong Province (Guangrao county had the highest HV carriage rate, at 16.67%), while the surveillance sites in southeastern Shandong Province had lower HV carriage rates(Pingdu city had the highest HV carriage rate, at 6.67%). Obviously, there were distinct differences in the spatial distributions of different HV-infected rodent species and HV carriage rates in Shandong Province.

**Table 3.** The results of laboratory testing of rodents captured in 2012-2015

| Rodents species | No. positive/no. tested (%) | Virus Type* |  |  |
| --- | --- | --- | --- | --- |
|  |  | HTNV | SEOV | Others |
| *R. norvegicus* | 93/908 (10.24) | 2 | 89 | 2 |
| *A. agrarius* | 17/269 (6.32) | 17 | 0 | 0 |
| *Mus musculus* | 1/373 (0.27) | 0 | 1 | 0 |
| *Sorex araneus* | 0/73 (0) | 0 | 0 | 0 |
| *Niviventer confucianus* | 0/25 (0) | 0 | 0 | 0 |
| *Miorotus fortis* | 0/1 (0) | 0 | 0 | 0 |
| *Cricetulus triton* | 0/31 (0) | 0 | 0 | 0 |
| *Rattus flavipectus* | 0/23 (0) | 0 | 0 | 0 |
| *Cricetulus barabensis* | 0/95 (0) | 0 | 0 | 0 |
| Total | 111/1,798 (6.17) | 19 | 90 | 2 |

HTNV= Hantaan virus; SEOV= Seoul virus.
\* All captured rodents were sent for laboratory testing and the HV carriage rates were also calculated. After the Blast process of partial L segments of the gene isolated from the positive samples, the genotypes were also figured out.

**Fig 4.**
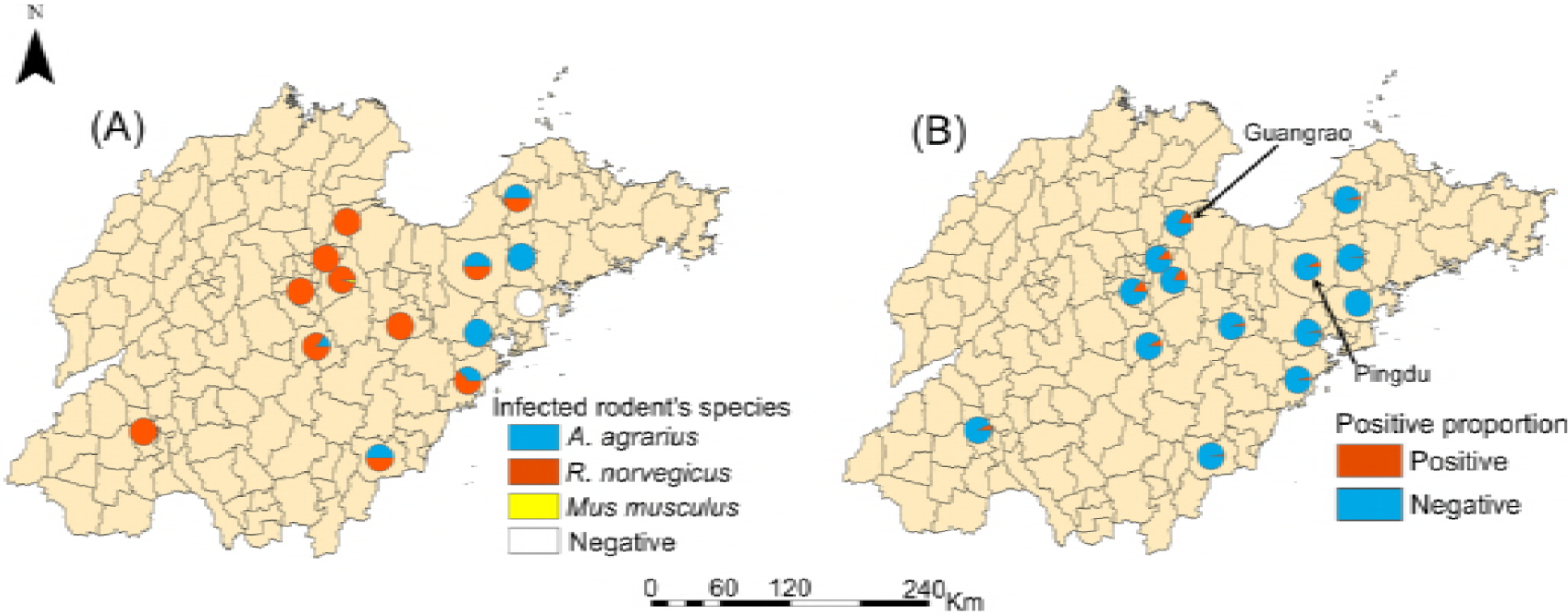
Spatial distribution of HV-infected rodents and the hantaviruses (HV) carriage rates in surveillance sites. (A) After laboratory testing, the species of HV-infected rodents in each surveillance sites were sorted out and displayed by different color. (B) The infection proportions of rodents captured in each surveillance site were also displayed. The map was visualized with ArcGIS 10.2 software (ESRI Inc., Redlands, CA, USA).

### Phylogenetic analysis

A total of 111 HV strains were isolated from the rodents captured in Shandong Province from 2012 to 2015. The Blast results using GenBank revealed that the numbers of SEOV and HTNV strains were 90 and 19, respectively, and two strains with novel genotypes were also found.

To construct a phylogenetic tree, partial L segment sequences from 44 HV strains were used: 29 strains from our samples and 15 strains from GenBank (which had been isolated in mainland China, the Korean Peninsula and Europe). As shown in Fig 5, the branches of the phylogenetic tree revealed that the HTNV strains isolated in this study consisted of two clades from a bigger branch. They shared a common ancestor with strains isolated in Hubei and Xi’an. The SEOV strains isolated in this study constituted one branch, except for one strain that was isolated in Jiaxiang County (in the southwest of Shandong Province). Almost all the SEOV strains were closely related to viruses isolated in Fujian, Beijing and Zhejiang. However, the SEOV strain isolated in Jiaxiang County was dissimilar to all other strains and formed a separate branch. In addition, the genetic diversity associated with the SEOV branches indicated that they were more homogeneous than the HTNV branches, which had a higher degree of genetic diversity.

**Fig 5.**
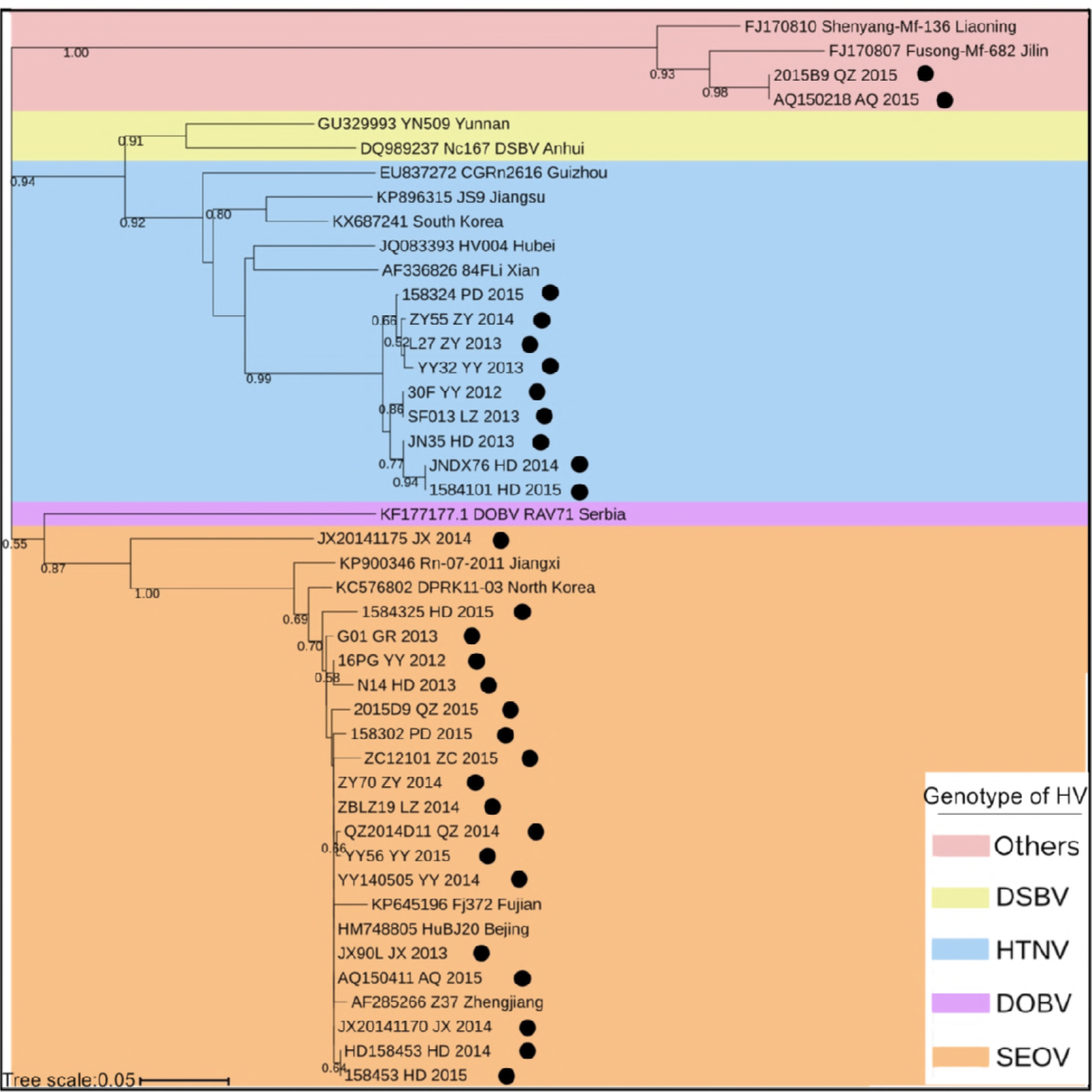
Phylogenetic tree of hantaviruses based on partial L segment sequences. The phylogenetic tree was constructed by partial L segment of 44 hantavirus (HV) sequences, including 29 sequences isolated in this study represented by black circles and 15 sequences downloaded from GenBank. The neighbor-joining (NJ) method and bootstrap testing (1000 replicates) were employed. Numbers above branches are proportions bootstrap support values for 1000 replicates and the scale bars indicate 0.05 substitutions per site.

We isolated two viruses with novel genotypes from *R. norvegicus* in Anqiu City and Qingzhou City that were different from the other strains. After the alignment analysis in GenBank, it was found that they were closely related to strains previously isolated in Fusong County and Shenyang City (northeastern China), forming a single branch in the phylogenetic tree. Previous studies on the complete S segments of strains isolated in Fusong County and Shenyang City have shown that they are closely related to a virus isolated in Vladivostok in the Russian Far East, forming a single branch. The researchers believed that they belong to the same HV genotype, the Vladivostok virus (VLAV). Therefore, our two novel viruses appear to be similar to the VLAV genotype [12,13].

### Analysis of HFRS epidemic foci

To more intuitively identify the current HFRS epidemic foci in Shandong Province, the distribution of HV genotypes in each surveillance site was mapped. Additionally, based on the two annual incidence peaks, we divided the 2012-2015 cases into three groups according to the month of their occurrence (Feb to May, June to Sept, and Oct to Jan) and mapped them with the 2012–2015 annual incidence rates on a single map. Finally, as shown in Fig 6, we summarized the 2012-2015 HFRS case spatial epidemiology, molecular epidemiology and rodent surveillance results in Shandong Province, indicating the epidemic foci during the study period.

The epidemic foci comprised two major foci, designated Regions a and b, which is located in southeastern and central Shandong Province, respectively, and a smaller one designated Region c in southwestern Shandong Province (S2 Table). Among them, Region a had the largest coverage area and the highest incidence rate, with most cases occurring in the autumn and winter, and the rodents carrying the virus were mainly *A. agrarius*. The viral genotypes in Region a were mainly HTNV, which constituted an *A. agrarius*-dominated mixed-type epidemic focus. The coverage and incidence rates of Region b were smaller than in Region a, the temporal distribution of cases in Region b was relatively uniform, and the infected rodents were mainly *R. norvegicus*. The viral genotypes in Region b were mainly SEOV, which constituted an *R. norvegicus*-dominated mixed-type epidemic focus. The coverage and incidence rates of Region c were much smaller than in Regions a and b. The cases in this area occurred mostly in late spring and early summer, and the species of infected rodents and viral genotypes were more similar to those of Region b, involving an *R. norvegicus*-dominated mixed-type epidemic area.

**Fig 6.**
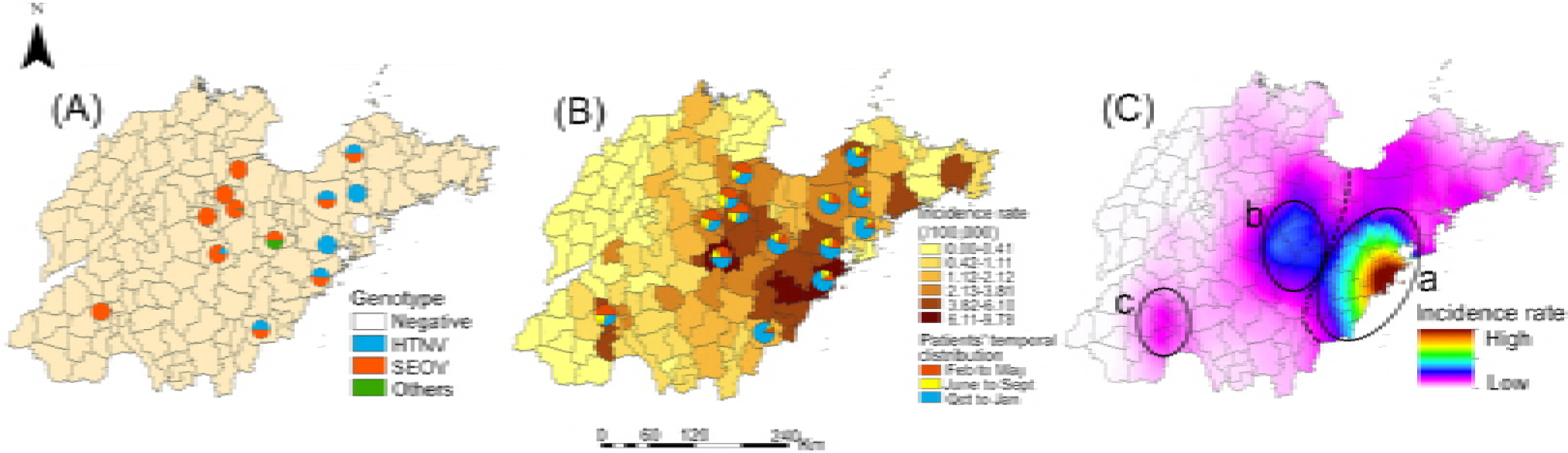
Status of epidemic foci of hemorrhagic fever with renal syndrome (HFRS) in Shandong Province from 2012 to 2015. (A) The Hantavirus (HV) genotypes isolated from each surveillance site were represented by different color. (B) The mean incidence rate of HFRS between 2012 and 2015 was calculated and overlapped with the temporal distribution of cases. The temporal distribution of cases was divided into three groups according to the month of their occurrence (Feb to May, June to Sept and Oct to Jan). (C) To get a clearer understanding of HFRS epidemic situation, the incidence rate of HFRS during the study was calculated by Kriging interpolation. Based on the results of spatial epidemiology, molecular epidemiology and rodent surveillance, the epidemic foci of HFRS in Shandong Province during the study period were summarized as the Regions a, b and c. The black dotted line represents the boundary between Regions a and b. The map was visualized with ArcGIS 10.2 software (ESRI Inc., Redlands, CA, USA).

## Discussion

As a natural epidemic disease, HFRS has a prominent position in the history of infectious disease prevention and control in Shandong Province. In Shandong Province, HFRS case peaked in the mid-1980s, then declined gradually and then became stable[6,14]. Based on the 2012-2015 HFRS data from Shandong Province, the characteristics of the disease, virus host and laboratory testing were analyzed in this study, and the current epidemic foci were identified.

Overall, the annual HFRS incidence rate exhibited small fluctuations during the study period, peaking in 2013, then decreasing year by year, and reaching its lowest point in 2015. There were still annual incidence peaks, which comprised a small peak in late spring to early summer and a big peak in autumn to winter. The small peak is associated with indoor-related infections linked to *R. norvegicus* breeding activity, while the big peak is mainly due to the increased contact between people and wild rats when they work in fields, and the migration of wild rats from fields to residential areas for foraging in the later period[14].

It is worth noting that 88% of all deaths occurred in September to January. Previous studies have suggested that the clinical symptoms of HFRS caused by HTNV are often more serious than those caused by SEOV[15]. Additionally, human immunity decreases in autumn and winter when it is cold. Therefore, the deaths may be the result of a combination of HTNV and seasonal factors. Additionally, male farmers represented the biggest HV-susceptible population because of their increased contact with rodents due to working in the fields and poor residential sanitation conditions[16]. Therefore, HFRS prevention and control in Shandong Province should continue to be focused on controlling the rodents population and strengthening the immunization and education interventions for farmers in key epidemic foci. To reduce the mortality rate, we should also pay more attention to the treatment of cases occurring in autumn and winter.

We found that although the HV carriage rate among rodents was lower in southeastern Shandong Province than in central Shandong Province, the epidemic situation and clinical symptoms of cases were worse. This suggests that more attention should be paid and more interventions should be undertaken in the southeastern region. Additionally, it’s uncommon to find HV in *Mus musculus* in Shandong Province before. Therefore, determining whether *Mus musculus* can carry HV and pass on the infection to humans should be a key task of future research.[17,18].

HTNV and SEOV were still the main HV genotypes in Shandong Province, but two points are worth noting. First, we isolated HTNV from *R. norvegicus* captured in the wild, indicating that HTNV from *A. agrarius* spilled over to *R. norvegicus*. Previous studies also showed that some HV isolates from *R. norvegicus* seemed to be HTNV[12,19,20]. We also found SEOV in *Mus musculus*. However, each HV genotype is associated primarily with a specific rodent host species that it coevolved with[17,21,22]. The phenomenon of viruses spilling over to non-traditional specific rodent hosts can allow them to find more suitable surroundings for a higher survival rate and more efficient infection and replication[23,24]. Second, we found two novel strains isolated from *R. norvegicus* in Anqiu City and Qingzhou City and they were closely related to strains from northeastern China, which reportedly belong to the VLAV genotype. This phenomenon of potential VLAV strains in Shandong Province has not previously been reported. Therefore, determining whether the viruses were transmitted from northeastern China, or simply involve indigenous genetic mutations, requires further research.

Since the first occurrence of HFRS in Shandong Province in the 1960s, it has undergone a change from *Rattus*-type epidemic foci centered on Jining City and *Apodemus*-type epidemic foci centered on Linyi City to mixture of them, then to an *R. norvegicus*-dominated mixed-type epidemic focus, and to a stable stage[7]. However, prior to the current study, we had no clear understanding of the HFRS epidemic foci in Shandong Province in the past decade. According to the results, the HFRS epidemic foci still involve *R. norvegicus*-dominated mixed-type epidemic foci.

In terms of the temporal distribution, the influence of Region a, which involved an *A. agrarius*-dominated mixed-type epidemic focus, on HFRS epidemic situation in Shandong Province mainly occurred in the autumn and winter. The coverage of Region a was more extensive than the coverage of Regions b and c, and the epidemic situation was also more serious. We inferred from previous studies that Region a is derived from the epidemic foci previously centered on Linyi City. Conversely, Regions b and c were more similar, involving *R. norvegicus*-dominated mixed-type epidemic foci. In Regions b and c, the temporal distribution of HFRS cases was more uniform and the epidemic situation was more moderate than in Region a. It was inferred that Region b is derived from the epidemic foci previously centered on Jining City, while Region c is the residue of the epidemic foci centered on Jining City[7].

Although the HV carriage rates among rodents in Regions b and c were higher than that associated with Region a, the severity of the epidemic situation was less severe, suggesting that the HV-susceptible population appears to be more susceptible to the HV strains in Region a (which mainly belong to the HTNV genotype). Interestingly, the distance between Regions b and a is much smaller than that between Regions b and c. Despite this, the epidemic characteristics of Region b are quite different from those of Region a, and more similar to those of Region c. Although previous studies have shown that HTNV (which was the main genotype in Region a, where the epidemic situation was more serious) is more harmful to humans than SEOV (which was the main genotype in Regions b and c), local geographic conditions, climate, HFRS prevention and control measures and safety awareness among susceptible populations are also important factors[15,25–29]. Therefore, we believe that a further study should be performed to explore the factors underlying our result.

Several limitations of this study should be noted. First, the HV genome can be divided into S, M and L segments. A comprehensive phylogenetic analysis of the three segments would give us a clearer understanding of the genetic diversity and evolutionary trajectory of HV. The data on S and M segments in this study were insufficient, so the phylogenetic analysis of HV may be biased. Second, there are many factors that can influence the distribution and severity of infectious disease, such as natural environmental factors, urbanization processes and local prevention efforts[29–31]. However, this research lacked information on these factors, so the HFRS spatio-temporal distribution analysis is inadequate.

In conclusion, the southeastern and central Shandong Province are current key HFRS epidemic foci dominated by *A. agrarius* and *R. norvegicus*, respectively. Although the overall epidemic intensity was lower than the previous intensity in Shandong Province, it was still relatively high compared to the rest of the country. The spatio-temporal distribution of HFRS within Shandong Province indicated different epidemic situations in different regions. Local departments still need to strengthen their prevention and control measures to reduce the hazards of HFRS.

## Supporting Information

**S1 Checklist.** STROBE Checklist.

**S2 Table.** The virus carriage rates of rodents captured in 14 surveillance sites, 2012-2015

**S3 Table.** The relationship between different partitions in Shandong Province in this study

## References

1. Zhang S, Wang S, Yin W, Liang M, Li J, et al. (2014) Epidemic characteristics of hemorrhagic fever with renal syndrome in China, 2006-2012. BMC Infect Dis. 14: 384.

2. Zou LX, Chen MJ, Sun L (2016) Haemorrhagic fever with renal syndrome: literature review and distribution analysis in China. Int J Infect Dis. 43: 95–100.

3. Kariwa H, Yoshimatsu K, Sawabe J, Yokota E, Arikawa J, et al. (1999) Genetic diversities of hantaviruses among rodents in Hokkaido, Japan and Far East Russia. Virus Res. 59: 219–228.

4. Wang T, Liu J, Zhou Y, Cui F, Huang Z, et al. (2016) Prevalence of hemorrhagic fever with renal syndrome in Yiyuan County, China, 2005-2014. BMC Infect Dis. 16: 69.

5. Wang S, Hang C, Wang H, Xie Y, Ma B (2002) Genotype and Clade Distribution of Hantaviruses in China. Chin J VIROLOGY.:211–216.

6. Fang LQ, Wang XJ, Liang S, Li YL, Song SX, et al. (2010) Spatiotemporal trends and climatic factors of hemorrhagic fever with renal syndrome epidemic in Shandong Province, China. PLoS Negl Trop Dis. 4: e789.

7. Liu YX, Wang ZQ, Guo J, Tang F, Sun XB, et al. (2013) Spatio-temporal evolution on geographic boundaries of HFRS endemic areas in Shandong Province, China. Biomed Environ Sci. 26: 972–978.

8. Wang H, Yoshimatsu K, Ebihara H, Ogino M, Araki K, et al. (2000) Genetic diversity of hantaviruses isolated in china and characterization of novel hantaviruses isolated from Niviventer confucianus and Rattus rattus. Virology. 278: 332–345.

9. Klempa B, Fichet-Calvet E, Lecompte E, Auste B, Aniskin V, et al. (2006) Hantavirus in African wood mouse, Guinea. Emerg Infect Dis. 12: 838–840.

10. Kim WK, No JS, Lee SH, Song DH, Lee D, et al. (2018) Multiplex PCR-Based Next-Generation Sequencing and Global Diversity of Seoul Virus in Humans and Rats. Emerg Infect Dis. 24: 249–257.

11. Zhang YZ, Zhang FX, Wang JB, Zhao ZW, Li MH, et al. (2009) Hantaviruses in rodents and humans, Inner Mongolia Autonomous Region, China. Emerg Infect Dis. 15: 885–891.

12. Zhang YZ, Zou Y, Fu ZF, Plyusnin A (2010) Hantavirus infections in humans and animals, China. Emerg Infect Dis. 16: 1195–1203.

13. Zou Y, Xiao QY, Dong X, Lv W, Zhang SP, et al. (2008) Genetic analysis of hantaviruses carried by reed voles Microtus fortis in China. Virus Res. 137: 122–128.

14. Kang D, Wang Z, Fu J, Yuan Q, Chen R, et al. (2007) Study on the incidence and spatiotemporal dynamic Variation of hemorrhagic fever with renal syndrome in Shandong province. Chin J Epidemiol. 28: 468–472.

15. Wu J, Wang DD, Li XL, de Vlas SJ, Yu YQ, et al. (2014) Increasing incidence of hemorrhagic fever with renal syndrome could be associated with livestock husbandry in Changchun, northeastern China. BMC Infect Dis. 14: 301.

16. Liang W, Gu X, Li X, Zhang K, Wu K, et al. (2018) Mapping the epidemic changes and risks of hemorrhagic fever with renal syndrome in Shaanxi Province, China, 2005-2016. Sci Rep. 8: 749.

17. Xiao H, Tong X, Huang R, Gao L, Hu S, et al. (2018) Landscape and rodent community composition are associated with risk of hemorrhagic fever with renal syndrome in two cities in China, 2006-2013. BMC Infect Dis. 18: 37.

18. Jiang F, Zhang Z, Dong L, Hao B, Xue Z, et al. (2016) Prevalence of hemorrhagic fever with renal syndrome in Qingdao City, China, 2010-2014. Sci Rep. 6: 36081.

19. Zou Y, Hu J, Wang ZX, Wang DM, Yu C, et al. (2008) Genetic characterization of hantaviruses isolated from Guizhou, China: evidence for spillover and reassortment in nature. J Med Virol. 80: 1033–1041.

20. Cao S, Ma J, Cheng C, Ju W, Wang Y (2016) Genetic characterization of hantaviruses isolated from rodents in the port cities of Heilongjiang, China, in 2014. BMC Vet Res. 12: 69.

21. Jiang H, Zheng X, Wang L, Du H, Wang P, Bai X (2017) Hantavirus infection: a global zoonotic challenge. Virol Sin. 32: 32–43.

22. Klein TA, Kim HC, Chong ST, Kim JA, Lee SY, et al. (2015) Hantaan virus surveillance targeting small mammals at nightmare range, a high elevation military training area, Gyeonggi Province, Republic of Korea. PLoS One. 10: e0118483.

23. Liu J. Genetic analysis of hantaviruses and their hosts in Hubei province. (2011), Wuhan: Wuhan University.

24. Wang W, Wang MR, Lin XD, Guo WP, Li MH, et al. (2013) Ongoing spillover of Hantaan and Gou hantaviruses from rodents is associated with hemorrhagic fever with renal syndrome (HFRS) in China. PLoS Negl Trop Dis. 7: e2484.

25. He X, Wang S, Huang X, Wang X (2013) Changes in age distribution of hemorrhagic fever with renal syndrome an implication of China’s expanded program of immunization. BMC Public Health. 13: 294.

26. Jiang H, Du H, Wang LM, Wang PZ, Bai XF (2016) Hemorrhagic Fever with Renal Syndrome: Pathogenesis and Clinical Picture. Front Cell Infect Microbiol. 6: 1.

27. Krüger DH, Schönrich G, Klempa B (2011) Human pathogenic hantaviruses and prevention of infection. Hum Vaccin. 7: 685–693.

28. Li S, Ren H, Hu W, Lu L, Xu X, et al. (2014) Spatiotemporal heterogeneity analysis of hemorrhagic fever with renal syndrome in China using geographically weighted regression models. Int J Environ Res Public Health. 11: 12129–12147.

29. Fang LQ, Goeijenbier M, Zuo SQ, Wang LP, Liang S, et al. (2015) The association between hantavirus infection and selenium deficiency in mainland China. Viruses. 7: 333–351.

30. Yan L, Fang LQ, Huang HG, Zhang LQ, Feng D, et al. (2007) Landscape elements and Hantaan virus-related hemorrhagic fever with renal syndrome, People’s Republic of China. Emerg Infect Dis. 13: 1301–1306.

31. Wei L, Qian Q, Wang ZQ, Glass GE, Song SX, et al. (2011) Using geographic information system-based ecologic niche models to forecast the risk of hantavirus infection in Shandong Province, China. Am J Trop Med Hyg. 84: 497–503.

